# Cross-species Transmission and Host Adaptation of Feline Leukaemia Virus between Domestic Cats and the Wild Felid *Leopardus guigna*

**DOI:** 10.64898/2025.12.17.694864

**Authors:** Cristobal Castillo-Aliaga, Irene Sacristán, Camila J. Stuardo, Constanza Napolitano, Ezequiel Hidalgo-Hermoso, Rachael E Tarlinton

**Affiliations:** School of Veterinary Medicine and Science, University of Nottingham, Sutton Bonington, UK; Epidemiology and Environmental Health Group, Department of Infectious Animal Diseases and Global Health, Animal Health Research Centre, National Centre Institute for Agriculture and Food Research and Technology, Spanish National Research Council (CISA-INIA-CSIC), Madrid, Spain; Departamento de Ciencias Biológicas y Biodiversidad, Universidad de Los Lagos, Av. Fuchslocher 1305, Osorno, 5311157, Chile; Institute of Ecology and Biodiversity, Victoria 631, Concepción, 4070374, Chile; Cape Horn International Center, O’Higgins 310, Puerto Williams, 6350000, Chile; Centro de Conservación de la Biodiversidad Chiloe Silvestre

**Keywords:** Feline Leukemia Virus, Domestic cats, NGS, Wild Felid, Illumina

## Abstract

Feline leukaemia virus (FeLV) is widespread in domestic cats and frequently spills over into wild felid populations, causing severe outbreaks in non-domestic felid populations. Domestic cats carry both endogenous FeLV (enFeLV) and exogenous variants of FeLV (exFeLV), which can recombine to generate novel variants with alternate receptor usage. In contrast, most non-domestic felids, including the guigna are lack enFeLV

This study applied amplification of the FeLV envelope gene combined with Illumina sequencing to characterise FeLV envelope gene diversity and transmission dynamics in guignas (Leopardus guigna), a small wild felid native to Chile and Argentina. Seven PCR amplicons from five free ranging guignas were sequenced.

Illumina contigs demonstrated that infections originated from Chilean domestic cats; however, a distinct guigna-specific cluster with unique sequence variations was identified, suggesting that FeLV transmission also occurs among guignas independent of domestic cats. Phylogenetic analysis showed that all guigna sequences formed a Chile-specific FeLV clade, closely related to domestic cat viruses but clearly separated from international FeLV-A strains.

## 3. Introduction

Feline leukaemia virus (FeLV) is an enveloped retrovirus that belongs to the family *Retroviridae*, genus *Gammaretrovirus* (Coffin et al. 2021). The FeLV genome consists of two copies of a single positive sense RNA around 8.4 kb in length. It encompasses three genes: *gag* (capsid proteins), *pol* (reverse transcriptase enzymes, protease, and integrase), and *env* (envelope proteins). These are flanked, 5’ and 3’, by two identical untranslated regulatory sequences called LTRs (long terminal repeats). During infection, the RNA genome is copied into a DNA genome by the reverse transcriptase enzyme and is inserted as a provirus into the host genome, leading to lifelong infection and facilitates recombination events to contribute to the FeLV diversity (Willett and Hosie 2013).

FeLV has a world-wide distribution in domestic cat populations and is one of the most important pathogens for their health (Hartmann and Hofmann-Lehmann 2020). The virus can replicate in many tissues, the most important being lymphoid tissues, salivary glands, and intestinal tracts (Helfer-Hungerbuehler et al. 2015). It causes clinical signs such as immune and bone marrow suppression, lymphadenopathy, lymphoma, leukaemia, oral lesions and respiratory diseases (Hartmann 2012). The virus is excreted in saliva, urine, faeces, milk and nasal secretions (Torres, Mathiason, and Hoover 2005). Social activities like grooming, sharing food and water dishes, nursing and bites during fights are the main transmission routes. It can also be transmitted via blood contact (blood transfusion or contaminated instruments) or sharing of litter trays (Little et al. 2020). Another important route of transmission is vertically, where the queen cat spreads virus to her neonatal kittens, either transplacentally, during parturition or during nursing of kittens (Hardy et al. 1976). FeLV also has importance in wild felid health and conservation causing severe disease and mortality in genetically bottle-necked populations such as the Iberian lynx (*Lynx pardinus*) (Meli et al. 2010a) and is notably harmful in the Florida panther (*Puma concolor coryi*) (Chiu et al. 2019, Cunningham et al. 2008). Although, the virus can persist in wild species, domestic cats have been the infection source for non-domestic animals in the majority of cases, including Iberian lynx (Meli et al. 2010c), Florida panthers (Brown et al. 2008a) and both pumas and bobcats (Petch et al. 2022). Thus, FeLV represents a critical pathogen at the domestic–wildlife interface.

On the other hand, endogenous FeLV (enFeLV) are retroviruses that have integrated over millions of years into the cat genome. EnFeLV were acquired during the divergence of the Felidae and are present in domestic cat chromosomes and their close relatives in the *Felis* genus. They are transmitted from parents to descendants through chromosomes (Polani et al. 2010). The presence of enFeLV can result in recombination events with exogenous FeLV (exFeLV) and produce FeLV-B, a more virulent variant with altered cellular tropism (Willett and Hosie 2013). Because non-domestic felids (except *Felis silvestris*) lack enFeLV, their FeLV infections provide a unique opportunity to study exogenous viral evolution without interference from endogenous elements (Polani et al. 2010).

In domestic cats, FeLV-A is the most widespread group that can be horizontally transmitted and is often described as the least pathogenic subtype (Ahmad and Levy 2010, Biezus et al. 2023). All other groups arise de novo from FeLV-A mutation or by recombination with endogenous elements (Hartmann and Hofmann-Lehmann 2020). FeLV-B is the second most common subgroup, being detected in roughly half of all FeLV-A cases (Powers et al. 2018). Overall, FeLV-B is associated with worse clinical symptomatology and prognosis (Ahmad and Levy 2010). Although FeLV-B has historically been described as unable to be horizontally transmitted without FeLV-A, studies in domestic cats may suggest that FeLV-B transmission may occur (Erbeck et al. 2021, Stewart et al. 2013). In particular a more robust case was documented in a puma infected by FeLV-B, without FeLV-A, these subgroup dynamics highlight the evolutionary flexibility of FeLV and the importance of studying its genetic variability across domestic and non-domestic species (Chiu et al. 2019).

The majority of these variations occur in the *env* gene, which is the binding point between the virus and host cell (Cano-Ortiz et al. 2022). The *env* gene is composed of the Surface protein (SU) and Transmembrane protein (TM). SU includes three regions from 5’ to 3: Receptor binding domain (RBD), Proline rich region (PRR) and C-Domain. The RBD and C-Domain are involved in cellular tropism, therefore changes in these regions can modify cell receptor usage and so change clinical progression (Faix et al. 2002, Cano-Ortiz et al. 2022). More specifically, there are three main Variable Regions (VRA, VRB, VRC) identified, along with specific amino acid changes which are associated with changes in receptor usage (Boomer et al. 1997).

FeLV epidemiology in domestic cats is related to multiple factors, developed countries generally have a controlled situation with FeLV, showing prevalence in Europe between 0.7%-5.5% (Studer et al. 2019), and 3.1% FeLV prevalence in North America (Burling et al. 2017). Conversely, the situation in South America, is highly variable ranging from 3% in north-eastern Brazil (Lacerda et al. 2017) to 59.44% in Colombia (Ortega, C. et al. 2020), whilst in Chile, the prevalence has been described as between 20.2%-33% in rural areas (Mora et al. 2015, Sacristán et al. 2021a), and between 14% to 54.5% in cities (Castillo-Aliaga et al. 2024, Choi et al. 2024). This marked heterogeneity reflects differences in cat management, human population density, and veterinary care access across the continent and within Chile.

In South American non-domestic species, molecular detection of FeLV has been reported in guignas (*Leopardus guigna*) from Chile (Mora et al. 2015, Sacristán et al. 2021a), ocelots (*Leopardus pardalis*) from Ecuador (Villalba-Briones et al. 2022), and multiple species from Brazil, including ocelots, oncillas (*Leopardus tigrinus*), jaguarundis (*Puma yagouaroundi*) (Filoni et al. 2017), and jaguars (*Panthera onca*) (Guimaraes et al. 2009, Silva et al. 2016). In North America, FeLV has been associated with clinical signs and pathological findings in a captive bobcat (*Lynx rufus*) in the USA (Sleeman et al. 2001), as well as outbreaks in Florida panthers (*Puma concolor coryi*) and some cases in bobcats (Petch et al. 2022, Chiu et al. 2019, Brown et al. 2008, Cunningham et al. 2008). These repeated spillover events have demonstrated the capacity of FeLV to cross species barriers and establish sustained transmission in susceptible wild felids.

The Florida panther and Iberian Lynx (*Lynx pardinus*) have been the most emblematic examples of FeLV impact in non-domestic felids. The worst consequences of FeLV introduction in these species has been the exacerbation of the consequences of genetic bottlenecks, causing population size reductions in already vulnerable populations (Brown et al. 2008, Cunningham et al. 2008, Palomares et al. 2010, Meli et al. 2010a, Chiu et al. 2019, Nájera et al. 2024). The most recent FeLV survey in North America demonstrated a prevalence of 3.12% in pumas, 0.47% of bobcats, and 6.25% in domestic cats. A total of 20 transmission events was inferred, three domestic cats to puma spillovers, three confirmed puma-to-puma transmissions, and 14 events likely to have originated from domestic cats (Petch et al. 2022). The Iberian lynx has also been severely affected by FeLV (López et al. 2014). Between 2003-2007, at the beginning of the outbreak, 21% of animals tested positive for FeLV, and 6 individuals died from FeLV-related disease (Meli et al. 2009). As a consequence, intensive management strategies have been implemented in both species, including vaccination campaigns, isolation of progressively infected animals, and translocations to decrease the inbreeding rates and disease risk (Nájera et al. 2021, 2024, Palomares et al. 2010, Cunningham et al. 2008, Chiu et al. 2019b). After more than a decade of intervention, the epidemiological situation has shifted from acute mortality (Cunningham et al. 2008, Brown et al. 2008, Luaces et al. 2008, Meli et al. 2010) to more controlled scenarios that allow continuous surveillance. This has deepened the understanding of FeLV dynamics in endangered species. However, changes in population density and reduced habitat availability have led to greater overlapping areas and higher tolerance of intra-species interactions, facilitating for example puma-to-puma transmission (Cunningham et al. 2008, Petch et al. 2022, Chiu et al. 2019, Kraberger et al. 2020). Despite this, domestic cats remain the most likely source of initial infection (Meli et al. 2009, Chiu et al. 2019, Petch et al. 2022), although there is a risk that some viral variants may become more adapted to different species such as FeLV-B in pumas (Chiu et al. 2019). Additionally, reports of enteritis associated with FeLV have been described (Filoni et al. 2017, Gregor et al. 2024), expanding the traditional profile of FeLV as the cause of tumoral and immunosuppressive disease, expanding the knowledge about how FeLV can severely affect endangered felids and highlight the importance of early detection and molecular monitoring.

The guigna has been classified as Vulnerable for over a decade, with a continually decreasing population trend due to severe fragmentation of its habitat. It has recently been downlisted to Least Concern due to better information available on the species, though the same threats are still present across it’s range (Napolitano et al. 2015, Gálvez et al. 2025). Like other felids, guignas are solitary animals, except during mating (Sanderson, Sunquist, and W. Iriarte 2002). Reduced genetic diversity has been correlated with human-dominated landscapes (Napolitano et al. 2015), as well as pathogen infections associated with the presence of humans and domestic animals (Sacristán et al. 2021b, 2021a, Ortega, R. et al. 2020). In contrast to the managed FeLV outbreaks in the USA and Spain, guignas in Chile have shown even higher FeLV prevalence (>20%) than observed in Florida panthers or Iberian lynxes (Mora et al. 2015, Sacristán et al. 2021). This could reflect increased interactions with domestic cats due to habitat disruption, or more frequent guigna to-guigna contact. Moreover, FeLV remains uncontrolled among domestic cats in Chile (Castillo Aliaga et al. 2024), similarly to other Latin American countries (Ortega, C. et al. 2020, Acevedo et al. 2020, Santana, Pozo, and Castañeda 2022). Despite the high prevalence and conservation relevance of FeLV in guignas, no studies to date have characterised full env gene diversity, intra-host variation, or phylogenetic structure using next-generation sequencing.

Based on this information, we conducted a molecular survey for FeLV infection in free ranging guignas using targeted *env* amplification followed by Illumina sequencing to characterise viral diversity and transmission patterns. Our objective was to determine the genetic diversity of FeLV circulating in guignas and identify the likely source of infection by comparison with domestic cat sequences. We hypothesize that FeLV infections in guignas originate primarily from domestic cats, although subsequent guigna-guigna transmission may been also occurring, leading to distinct viral variants and unique intra-host mutation patterns. Therefore, we expected to find i) phylogenetic clustering of guigna FeLV sequences with Chilean domestic cat FeLV-A strains, and ii) guigna-associated cluster/s with nucleotide and amino acid substitutions suggestive of viral divergence within the species

## 4. Methods

### Samples and End-point PCR

Free-range guigna samples were sourced from previous studies with permission from the Chilean Agriculture and Livestock Service (Capture permits 814/13 2008, 109/9 2009, 1220/22 2010,1708/26 2010, 7624/2015, 2288/2016, 2185/2017, 4072/2018) and approved by the Animal Ethics Committee (Institute of Ecology and Biodiversity, resolution 20 November 2015). Guigna samples were captured from rural areas of the Southern area of Chile (Figure 1), these were: Puerto Cisnes (44°31′25.68″S 72°34′12.18″W), Paine (33°52′21.71″S 70°55′56.30″W), Hualañe (34°53′21.86″S 71°46′21.92″W), and two locations in Ancud (41°50′25.21″S 73°36′20.98″W; 41°51′27.73″S 73°35′34.03″W). South America map was generated in Rstudio using “rnaturalearth” (Massicotte and South 2023) ggmap (Kahle and Wickham 2013), and Google maps (2025).

**Figure 1.**
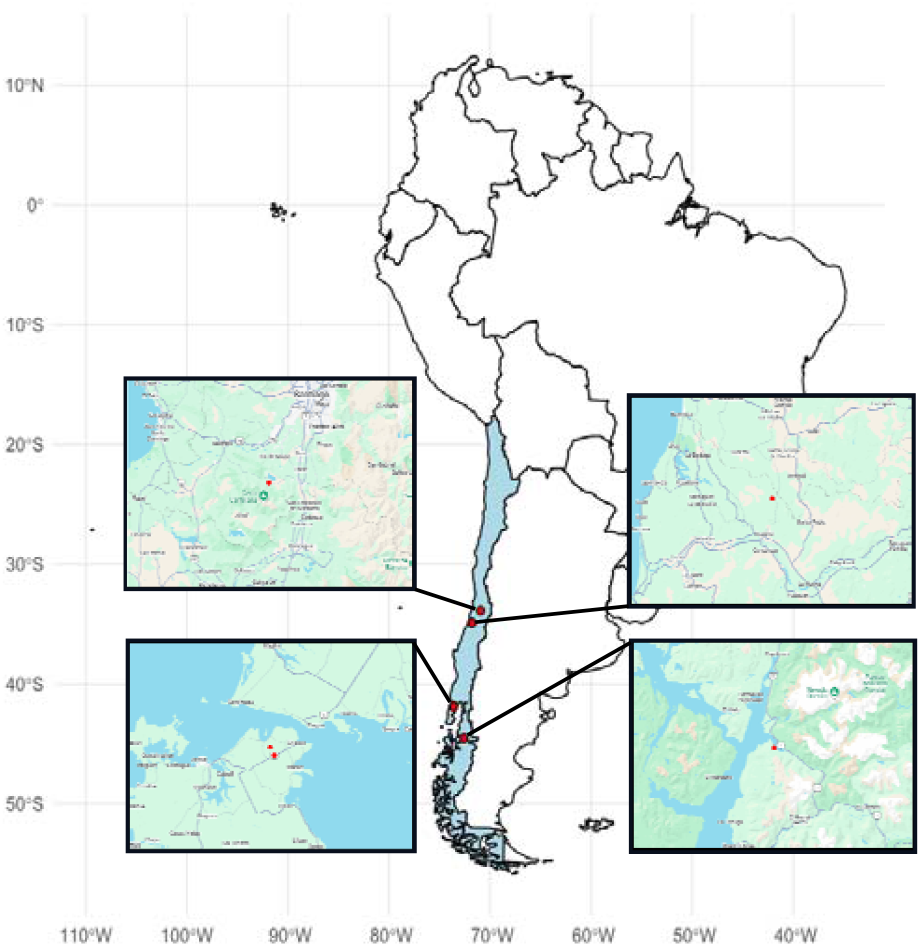
Map of South America and the geographical location of Chile (in light blue). Small circles indicate the sites where guignas were sampled. The five individuals were collected from four locations: Hualañe, Paine, Puerto Cisne, and two from Ancud

The samples from free ranging guigna were processed at the Universidad de Los Lagos, Osorno, Chile. In all cases, DNA extraction was performed using the commercial QIAGEN DNeasy Blood and Tissue kit following manufacturer’s instructions (Qiagen, Valencia, California, USA).

End-point PCR using two primer pairs following methods from Erbeck *et al*, (2020) was used to amplify the envelope gene hypervariable region and 3’LTR region from FeLV-A and FeLV-B. PCR was performed using GoTaq® Long PCR Master mix (Promega) and each reaction consisted of: 12.5µl of GoTaq® Long PCR Master mix; 0.5µl (10µM) Forward primer; 0.5µl (10µM) Reverse primer; 9.5µl of Nuclease-Free water; 2µl of template DNA to reach a final volume of 25µl per reaction.

The same protocol was used for each pair of primers. Thermo-cycler conditions used followed the Erbeck *et al*., (2020) protocol with some modifications according to the polymerase requirements. The protocol was: initial denaturation at 94ºC for 2 min followed by 45 cycles of denaturation at 95ºC for 30 s; annealing at 63 ºC for 30 s; extension at 72ºC for 2 mins and a final extension at 72ºC at 2 mins. All PCR runs included extraction blanks and non-template controls to monitor contamination. The thermo-cycler conditions were tested with a gradient PCR to select the optimal annealing temperature, and every set of PCR reactions included a negative control with no template (DNA free water) and a positive control (domestic cat FeLV-A DNA). The PCR products were run by electrophoresis on a 1% agarose TAE (Fisher scientific®) gel at 80V, 400 mA for 60 minutes. DNA purification used the Nucleospin® extract II kit (Macherey-Nagel®), according to the manufacturer’s instructions.

DNA quantification was performed with a Qubit 4 Fluorometer (Thermo Fisher Scientific®). FeLV-A and FeLV-B amplicons of high enough quality and quantity were selected for Illumina sequencing.

### Genome sequencing and Read Processing

Amplicons from *env* genes were amplified and sequenced by the Illumina platform (HiSeq4000) with 300-bp paired-end distances using a coverage of 30x by Novogene Europe, Cambridge, United Kingdom. Library preparation followed Novogene standard protocol for amplicon sequencing.

Illumina reads were trimmed, and adaptors removed using FastP v0.23.1 (Chen et al. 2018) with default stringency parameters. De Novo Assembly from Illumina reads was performed using MEGAHIT (Li et al. 2015). The contigs generated by MEGAHIT were mapped to FeLV-A FAIDS (M18247) using Bowtie2 (Langmead and Salzberg 2012). All contigs were extracted and imported to Geneious v2023.0.4 (Biomatters Inc., Newark, NJ) to annotate and continue phylogenetic analysis. The dataset was constructed in Geneious using all available complete *env* gene FeLV related sequences and were annotated and aligned with MAFFT (Katoh and Standley 2013).

Data are available in GenBank under BioProject number PRJNA1328684. The corresponding SRA files are available using the following accession numbers: Lpgui1a (SRR35412604); Lpgui1b (SRR35412603); Lpgui2a (SRR35412602); Lpgui2b (SRR35412601); Lpgui3 (SRR35412600); Lpgui4 (SRR35412599); and Lpgui5 (SRR35412598).

The alignment was sent to the IQTREE web server (Trifinopoulos et al. 2016), where a maximum likelihood tree was inferred using the ModelFinder (Kalyaanamoorthy et al. 2017) to select the best-fit model nucleotide substitution. The model chosen was general time reversible (TIM2+F+G4), branch support analysis was ultrafast with 1,000 bootstrap replicates (Hoang et al. 2018). EnFeLV was used as an outgroup to root the tree, and the phylogenetic tree was visualized in FigTree v1.4.4 (Rambaut 2018).

### Intra-host Variation Analysis and Recombination Analysis

The intra-host single nucleotide variation (iSNV) analysis was done using BWA-MEM (Li 2013) and iVar (Grubaugh et al. 2019). Whole libraries were mapped to the FeLV-A sequence (M18247) using BWA-MEM and consensus sequences were generated using the iVar consensus command for each library. All reference sequences generated by iVar were cut at nucleotide 66 (M18247), to initiate at the same nucleotide position. The output was used to call single nucleotide variants and indels in iVar. The minimum base quality parameter was 10 (Q>10) and only variants meeting above the frequency cutoff 1% (p < 0.05) were retained.

## 5. Results

The *env* gene was successfully sequenced from 5 guignas, corresponding to 5 FeLV-A amplicons and 2 FeLV-B amplicons. All resulting contigs were approximately ∼1.9kb in length. The FeLV-A PCR amplicons matched the expected amplicon size (∼1.9kb) and did not show the insertions or deletions commonly found in other variants of FeLV.

Although FeLV-B specific primers yielded amplicons, they revealed only FeLV-A sequences with random nucleotides attached to the 5’-end. The FeLV-A contigs shared 97.2% to 100% nucleotide identity among themselves and 98.8-100% identity at amino acid level. When compared to FeLV sequences from Chilean domestic cats (Castillo-Aliaga et al. 2023), the guigna sequences showed 97.2-99% nucleotide identity and 95.6%-99.7% amino acid identity.

BLASTn analysis revealed that the guigna sequences shared 98.86% identity with FeLV-FAIDS strain (GenBank accession M18247.1), and 98.44% identity with FeLV-A_Fca2018 from the US (ON995432.1). The FeLV-B amplicons demonstrated a similar identity percentage, although with a reduced query coverage due to random nucleotides attached to the 5’ end.

Contigs assembled using MEGAHIT were aligned with FeLV-A sequences from domestic cats in Europe, the US, Japan, Brazil, and Chile, as well as with sequences from non-domestic felids such as pumas and Iberian lynx, when available. Guigna samples clustered together in a robust phylogenetic separation (>90% of bootstrap support), without distinction between FeLV-A and FeLV-B amplicons (Figure 2). This guigna clade is adjacent to the cluster of Chilean domestic cats and was positioned near a puma sequence from the US. All were located inside a bigger cluster that includes the FeLV-FAIDS (M18147.1), and Rickard (AF052723) reference strains.

**Figure 2.**
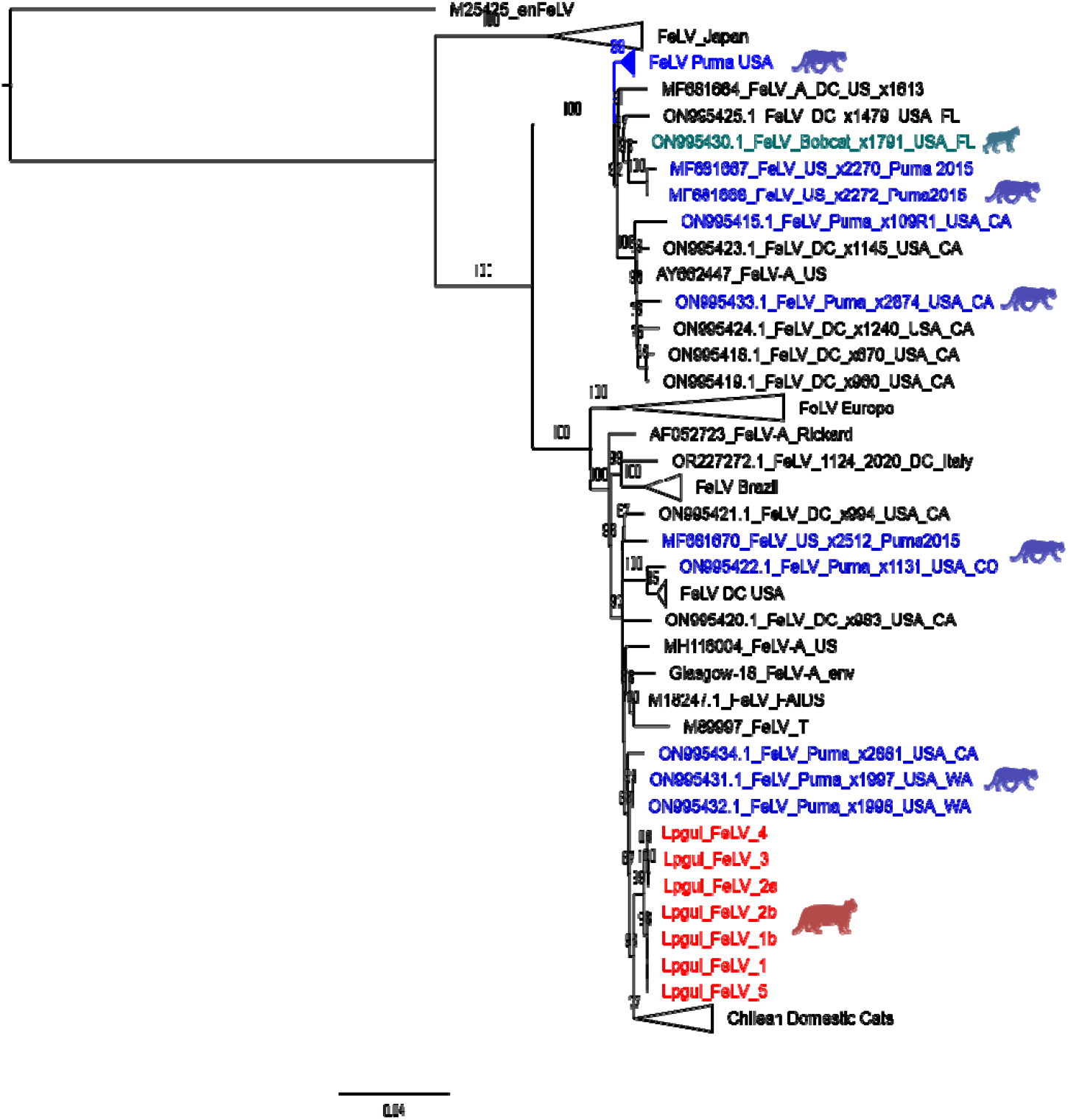
Phylogenetic tree of the FeLV env gene (nucleotide sequence) was constructed using 1,000 bootstrap approximations and rooted against enFeLV (M25425). The analysis includes only FeLV-A sequences from domestic and non-domestic felids, including all previously reported Chilean sequences. Guigna sequences are shown in red and form a distinct cluster adjacent to the Chilean domestic cat clade. Puma sequences are shown in purple, bobcat sequence is shown in turquoise. Guigna silhouette credit: Phylopic/Gabriela Palomo-Munoz CC BY 4.0

The iVar analysis identified a total of 96 intra-host single nucleotide variants (iSNVs), corresponding to 43 unique nucleotide positions across the 7 amplicons obtained from 5 guigna individuals (Figure 3). Overall iSNV counts per animal ranged from 2 to 52, indicating strong inter-individual heterogeneity. Among them, Lpgui_B7 and Lpgui_B123 showed the highest number of iSNVs with substantial variation also observed in sample Lgui_A169. In contrast, Lpgui_A128 presented only two iSNVs, while Lpgui_152 showed 8 iSNVs but with comparatively higher variant frequencies compared with the other animals (Figure 4).

**Figure 3.**
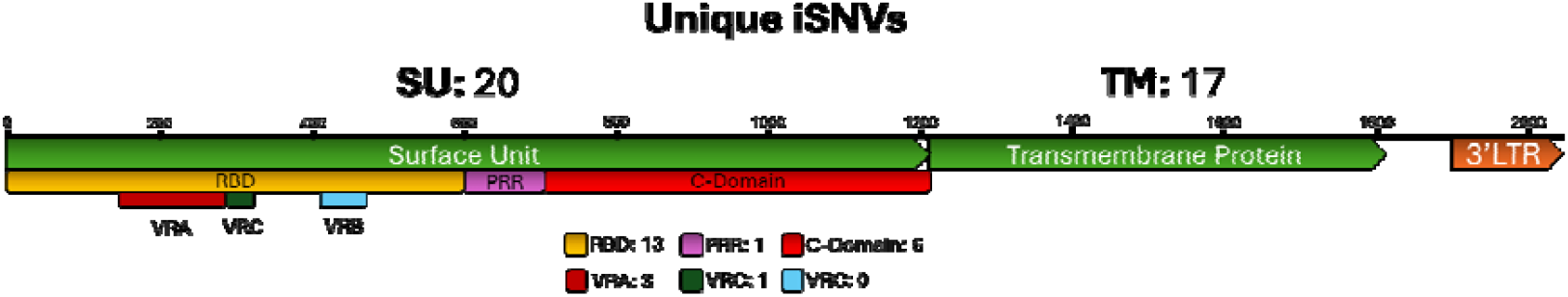
Genome structure amplification scheme showing unique iSNVs across the 7 amplicons from 5 guignas. Twenty events occurred in Surface Unit and 17 occurred in the Transmembrane Protein.

**Figure 4.**
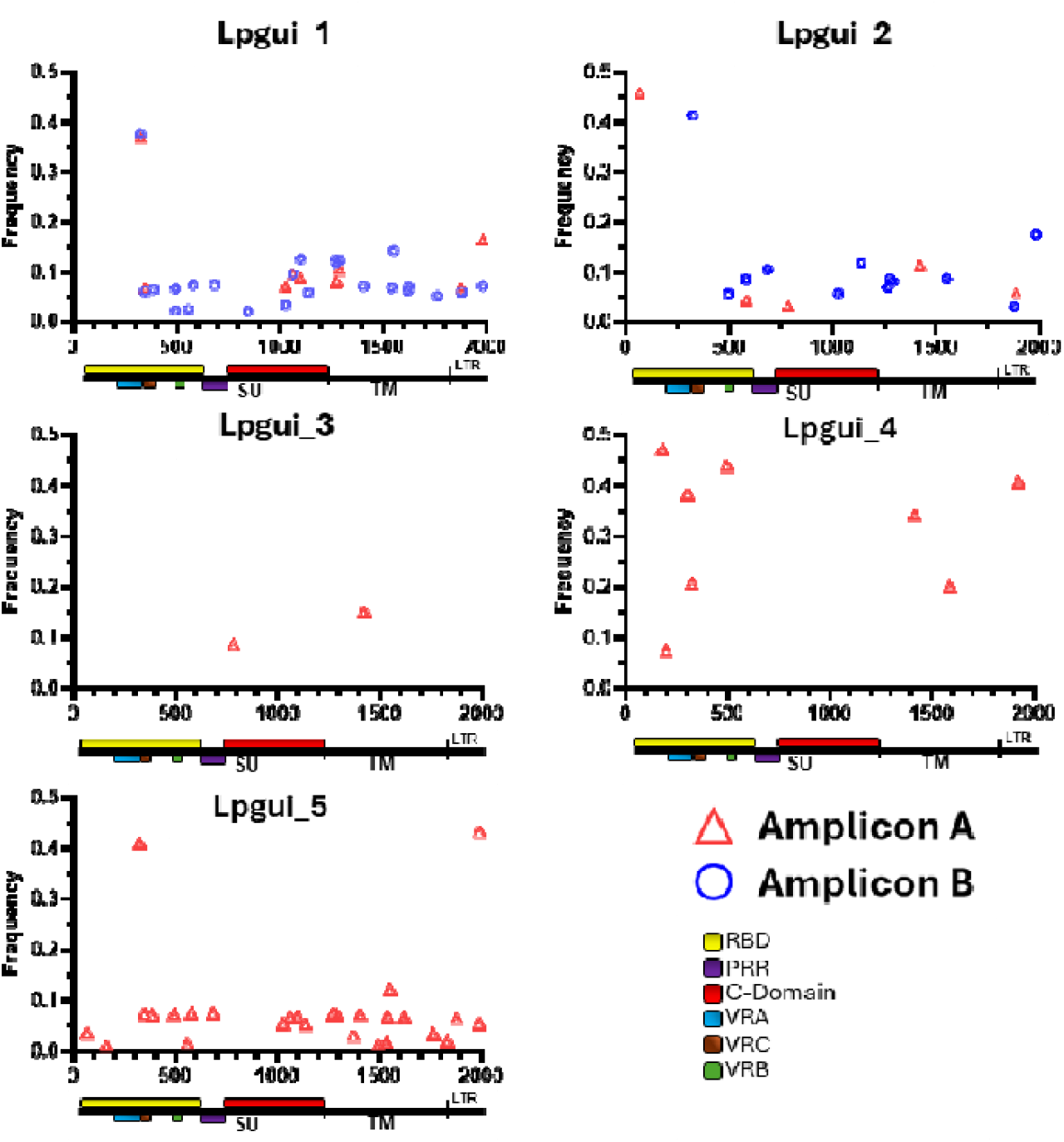
**iSNV sites detected in env gene amplicons from Illumina sequencing. iVar software was used to generate a consensus sequences per library and to be used as reference sequence (∼1.8kb). Blue circles represent iSNV obtained using primers to amplify FeLV-B and red triangles represent iSNV obtained with FeLV-A primers. Each graph represents a different animal.**

**Figure 5.**
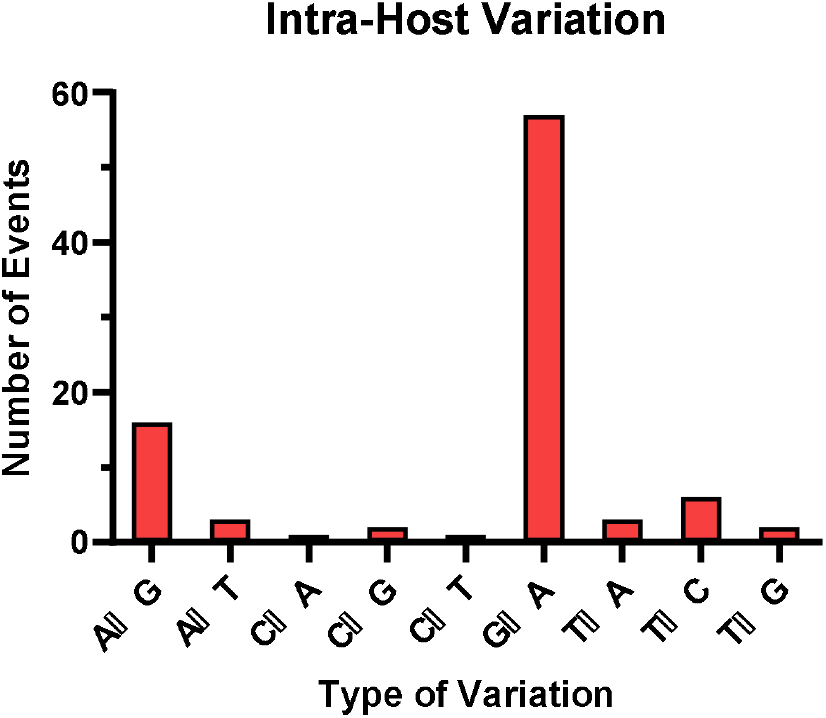
Bar graph of nucleotide variation in sequences obtained from Illumina sequencing for guigna samples.

In terms of genomic distribution, the majority of iSNVs (50) were in the in SU region, followed by 34 positions within TM region. Only 7 iSNVs were identified in the LTR region. The proportion of TM mutations (35.4%) was higher than typically reported in domestic cats, where TM is comparatively conserved. Within the SU domain, 31 iSNVs were identified in the RBD, 3 in the PRR, and 16 in the C-domain. As expected, the RBD was the most variable segment of the *env* gene.

Analysis of the type of iSNVs showed that G→A transitions were the most common, with a total of 57 events (62.6%). Other types of substitutions included 16 A→G (17.6%), 3 A→T and 3 T→A (3.3%), 2 C→G (2.2%), C→A and C→T had 1 event each (1.1%), 6 events of T→C 6 (6.6%), and 2 T→G (2.2%) (Figure 4).

Out of the total 96 iSNVs, 84 were located within coding regions, with 20 synonymous and 64 non-synonymous mutations. These non-synonymous variants were detected at different frequencies across samples, involving 26 distinct amino acid positions. Of these, 15 were G→A transitions, with 13 of them strongly suggestive of APOBEC-mediation, based on typical APOBEC motifs (AGG or GGG). Only one was less likely to be an APOBEC substitution (GAA to AAA).

Non-synonymous iSNVs were found in both the SU and TM regions, affecting 8 positions in SU and 7 in TM. The most frequently substituted amino acid was glycine replaced in 8 positions by Glutamic acid, Arginine or Serine. One notable iSNV at position 1620nt introduced a premature stop codon, potentially truncating the envelope protein and impacting viral function.

Based on the *de novo* assembled contigs, guigna samples were more conserved at the amino-acid level compared to Chilean domestic cat sequences. Four notable amino acid variations were identified differing between the Chilean domestic cats and other reference sequences (Table 1). At position 59 of the *env* gene, within the VRA, guigna sequences showed a serine (Ser), whereas Chilean cats presented either aspartic acid (Asp) or asparagine (Asn) at the same position. At position 60, three guigna contigs exhibited Ser, while 4 contigs showed proline (Pro), in contrast to domestic cats which consistently had Pro at this position. Afterwards, at position 71 all Chilean domestic cats and guignas, showed aspartic acid (Asp) compared with reference sequences. Position 165, however showed only variation in guignas, carrying either arginine (Arg) or lysine (Lys). Notably, position 249 consistently showed leucine (Leu) across all Chilean sequences (domestic cats and guignas), unique to the Chilean cluster and not identified in other sequences from GenBank.

**Table 1.** Amino acid variation identified across guigna and domestic cats.

| **Amino Acid position** | **59** | **60** | **71** | **165** | **249** |
| --- | --- | --- | --- | --- | --- |
| **DC_Chile** | D | P | D | R/K | L |
| **Lpgui_1a** | S | P | D | R | L |
| **Lpgui_1b** | S | P | D | R | L |
| **Lpgui_2a** | S | P | D | R | L |
| **Lpgui_2b** | S | S | D | K | L |
| **Lpgui_5** | S | S | D | K | L |
| **Lpgui_4** | S | S | D | K | L |
| **Lpgui_3** | S | S | D | K | L |
| **FAIDS** (M18247) | S | P | S | K | P |
| **Rickard** (NC_001940) | D | P | S | R | P |
| **Glasgow** (M12500) | N | P | S | R | P |
| **Iberian Lynx** (EU293175-94) | D | P | S | R | P |
| **Puma US** (ON995415-34) | D/S | P | S | R/K | P |

\*DC\_Chile: Chilean domestic cats from Castillo et al 2023

Several positions were recurrent across individuals: positions 325 and 1269 were present in 5 samples, while positions 494, 1027, 1269, 1276, 1291, and 1879 were found in 4 samples. Notably, the nucleotide common variants identified in the de novo assembly were also detected in other samples at lower frequencies, confirming common variant sites within the population.

For example, the amino acid substitution at position 61 was observed in sample A152 at a frequency of 47%, and in other samples at lower frequences (<10%). Similarly, that at position 169 was found at 43% in A152, and at 7% in both B123 and A169. Overall, the frequency of iSNVs was between 47% to 1% of total reads.

## 6. Discussion

The findings described in this study extend previous knowledge regarding the FeLV situation in guignas. Our results confirm that domestic cats are the primary source of FeLV infection for guignas with very close clustering of Chilean domestic cat and guigna sequences indicating region specific circulation of a Chilean FeLV-A lineage. We also identified a distinct guigna-specific FeLV cluster, characterized by unique mutations not observed in domestic cats. It is possible that very rapid adaption of virus to guigna is occurring, as occurs for avian flu isolates in mammals (Rather et al. 2025), but the complete separation of the guigna and domestic cat sequences in our study suggests that, while the virus originally arose from Chilean domestic cats, ongoing guigna-to-guigna transmission is now occurring. The receptor usage strongly influences host range and pathogenicity, in consequence, *env* gene sequencing is essential for understanding these cross-species transmission events.

Previous studies assessing infection prevalence across large regions of Chile have relied on 3’LTR diagnostic PCR (Mora et al. 2015, Sacristán et al. 2021a). This method is more sensitive considering that it is a smaller amplicon (211bp), and it is also highly conserved, decreasing false negative results (Miyazawa and Jarrett 1997). These characteristics make it suitable for prevalence calculations. However, analysis of the envelope gene, which is a hypervariable gene, provides additional information about the genetic dynamics of the virus (Watanabe et al. 2013, Petch et al. 2022, Chiu et al. 2019), including receptor usage for cell entry and potential clinical progression (Rohn et al. 1994, Cano-Ortiz et al. 2022), although the high similarity and recombination events between enFeLV and FeLV-A reduce the specificity of these primers, as we observed with our FeLV-B primers amplifying FeLV-A amplicons, and as previously described (Watanabe et al. 2013, Gallina et al. 2024).

In the US, FeLV in pumas has demonstrated multiple introduction events from domestic cat to pumas, followed by puma-to-puma transmission (Chiu et al. 2019, Petch et al. 2022). Two distinct FeLV-A clades were identified, resembling the situation in Japan where multiple FeLV-A clusters coexist (Petch et al. 2022, Watanabe et al. 2013). The epidemiological situation in pumas and Iberian lynx differs markedly from the guigna context. Both Iberian lynx and North American pumas are apex predators, frequently preying on domestic cats (Lee et al. 2017, Nájera et al. 2019, Brown et al. 2008). In contrast, the guigna is the smallest wild felid in the American continent, half the size (2 kg) of a domestic cat, with less frequent direct interactions (Napolitano et al. 2014). Contact between domestic cats and guigna occurs mainly at forest edges near rural human settlements, where guigna may enter human areas hunting poultry (Schüttler et al. 2017, Zorondo-Rodríguez, Reyes-García, and Simonetti 2014), and/or rural free-roaming domestic cats can move away from their households up to 2.5 km into forested areas (López-Jara et al. 2021), potentially leading to aggressive domestic cat-guigna encounters (Mora et al. 2015). This behavioural context likely also explains the higher prevalence in male guigna and Iberian Lynx (Meli et al. 2010, Sacristán et al. 2021a, Nájera et al. 2024). Following infection, males can transmit the virus to females during mating, as guigna, like most felids, are solitary outside of mating periods (Sanderson, Sunquist, and W. Iriarte 2002).

The phylogenetic tree presented in this study incorporated sequences previously described for domestic cats in Chile (Castillo et al. 2023) including samples from animals from diverse geographic origins, two other wild felid species, and using three different sequencing methods. A distinct Chilean cluster was clearly identified when compared to reference sequences from Europe and the USA (Cano-Ortiz et al., 2022; Gallina et al., 2024; Petch et al., 2022). Within this cluster, all Chilean sequences form a terminal cluster with a specific subcluster for guigna sequences. Interestingly, none of the Chilean sequences were related to the Brazilian strains, consistent with observations from the LTR-*gag* PCR analysis, demonstrating the circulation of distinct FeLV-A strains across South America.

At a first overview, the absence of large deletions or insertions in *env* suggest these viruses may not be highly pathogenic variants such as FeLV-C, T, B strains (Rohn et al. 1994, Stewart et al. 2012). Among the Chilean sequences, two amino acid changes were uniquely identified in guignas at positions 60 and 61, within the VRA region, which is critical for receptor interaction (Rohn et al. 1994, Stewart et al. 2012, Cano-Ortiz et al. 2022). The presence of Ser at position 60 has been described previously in Iberian lynx (Geret et al. 2011), some pumas (Petch et al. 2022), and domestic cats with tumours (Rohn, J L et al. 1994). However, the Ser at position 61 has not been previously described. While the presence of amino acid combinations of Asp52 and Asn60Asp mutations are known to enhance receptor binding, viral entry and viral replication (Stewart et al. 2013, Miyake et al. 2016), the significance of Ser at this position is unknown. These unique variations may represent specific viral adaptations to guigna receptors, and comparative studies on the sequence and structure of guigna FeLV receptors could elucidate the virus infectivity and replication efficiency in this wild felid. Another notable amino acid substitution, Pro249Leu, was observed exclusively in all Chilean FeLV strains, whether from wild or domestic felids and appears to be a feature of the currently circulating Chilean strains.

Evidence from Iberian lynx suggest that FeLV infection in this species is typically in a regressive state, with viral replication largely controlled except in a few progressively infected individuals (Nájera et al. 2024). This may partly explain the challenges in amplifying longer viral fragments observed here and in previous studies (Sacristán et al. 2021a, Gregor et al. 2024). A similar situation was described in a lynx in Germany, after an attempt to sequence the whole genome of FeLV using NGS yielded only partial sequences, requiring re-amplification with the Sanger method to complete the genome sequencing. This sequence showed notable differences regarding published reports, but there is no evidence of FeLV in other lynxes from that area (Gregor et al. 2024). Studies recorded FeLV PCR-positive guignas showing very low frequency of clinical signs, no histopathological evidence of disease and being serology negative, all suggestive of regressive infection (Sacristán et al. 2021a). Although we cannot definitively conclude that guigna experience similar regressive infections, the low diversity of FeLV strains in our samples and the low variation within each host suggest limited active replication and few new introductions of virus into the guigna population.

A notable finding was the unusually high proportion of iSNVs in guignas that occurred in the TM region of the *env* gene, resulting in amino acid variations and even premature stop codons. In domestic cats, in contrast, variation is typically concentrated in the SU region, especially the RBD area, while the TM domain remains highly conserved even between FeLV-A, FeLV-B and other recombinants (Cano-Ortiz et al. 2022).

A plausible explanation for the unexpected higher ratio of TM iSNVs is pressure from the guigna APOBEC antiviral system. APOBEC3 enzymes preferentially deaminate cytosines in AGG and GGG motifs (Stavrou and Ross 2015), generating higher numbers of G→A hypermutations that produce non-infective virions (Terry et al. 2017). Although APOBEC has a stronger impact on FIV and FFV than on FeLV in domestic cats (Münk et al. 2008), its activity may still drive the unique variation patterns we observe in guignas. A previous study using the Roche 454 sequencing platform to sequence the SU gene in Iberian lynx infections, reported some effects of the APOBEC system. They identified 25 sites of variations in SU, nearly half of which (48%) led to nonsynonymous changes, being predominantly Trp→Stop-Codon and Gly→Arg. (Geret et al. 2011). By contrast, our analysis additionally included TM protein and also indicated Gly as a frequently replaced amino acid. We identified a higher level of nonsynonymous variations (62.6%) and a sole premature stop codon was identified within the TM protein. The Iberian lynx study sampled fewer animals and detected fewer iSNVs per animal, while in our dataset one guigna harboured only two iSNVs and another showed over 50 different positions. Therefore, individual and species effects may also influence the efficacy of the APOBEC3 system in FeLV infections.

Comparative phylogenetic analyses of felid APOBEC systems indicates that the domestic cat and puma systems are closely related, whereas the lynx APOBEC diverged slightly earlier (Münk et al. 2008). Specifically, puma and domestic cat lineages diverged ∼6.7Mya, followed by lynx lineage divergence ∼7.2Mya (Werdelin et al. 2010). The *Leopardus* lineage (including guigna) diverged over 8 Mya (Werdelin et al. 2010, Lescroart et al. 2023). This evolutionary distance could indicate that guigna APOBEC3 may interact differently with FeLV genomes compared to domestic cat APOBEC3. Although it cannot be directly compared, FIV from pumas was less effective at infecting domestic cats, because the virus was less adapted to a different species and in consequence was more susceptible to domestic cat APOBEC (Münk et al. 2008). These findings support the idea that FeLV from domestic cats may likewise be poorly adapted to guigna APOBEC3, resulting in increased viral hypermutation in guignas.

When our iSNVs were mapped against published FeLV sequences, most appeared to be random, but several matched mutations documented in other hosts. For instance, the G109R substitution was originally described in domestic cats with tumours (Rohn, J L et al. 1994); the V262I variation had been described in cats suffering tumours and in Brazilian cats (Rohn, J L et al. 1994, Cano-Ortiz et al. 2022). Lys343Glu has been described across a wide range of hosts, including cats from US, an Iberian Lynx, European cats, the Rickard reference strain, Pumas, Brazilian cats (Chandhasin, Coan, and Levy 2005, Geret et al. 2011, Chen et al. 1998, Petch et al. 2022, Cano-Ortiz et al. 2022); Ala469Thr and Ala513Thr substitutions have been previously observed in US pumas (Chiu et al. 2019); and the Glu542 Lys mutation has reported in a US domestic cat (Erbeck et al. 2021).

Only two other studies have used NGS to characterize FeLV in non-domestic felids, one case report and one using one of the first NGS methods, which limits direct comparisons with our data set (Geret et al. 2011, Gregor et al. 2024). Nevertheless, we have successfully described *env* variation in guignas using Illumina sequencing. Given the receptor usage strongly influences host range and pathogenicity, env sequencing is essential for understanding cross-species transmission events. Another key advantage was the absence of enFeLV in guigna genomes, which reduced the interference of incorrect amplification and bioinformatic artefacts that were observed in domestic cat analysis (Castillo-Aliaga et al. 2023). The Illumina method proved highly effective for variant detection and produced results broadly consistent with those from domestic cats using the same method.

Rapid human migration from urban areas into Chilean native forest is driving significant landscape changes (Gálvez et al. 2018). As contact between domestic cats and other non-domestic felids increases, pathogen spillover risk will continuously increase. Additionally, this study provides evidence that there is likely guigna to guigna transmission occurring and highlights the need for monitoring for clinical impacts in this vulnerable species. It is therefore critical to maintain ongoing surveillance of FeLV both in domestic cats and in guigna populations. Intensive management strategies have been successfully implemented in both North American pumas and Spanish lynxes, including vaccination campaigns, isolation of progressively infected animals, and translocations to decrease the inbreeding rates and disease risk (Nájera et al. 2021, 2024, Palomares et al. 2010, Chiu et al. 2019, Cunningham et al. 2008), which may provide guidelines to establish measure for disease control. The effectiveness of the commercial domestic cat FeLV vaccines in controlling diseases has been widely demonstrated in the Iberian lynx and North American puma species recovery programmes and will be useful to establish in potential future guigna programmes.

## 8. Author Contributions Statement

CC: performed bioinformatic, molecular analysis, and wrote the manuscript. CN: conducted animal sampling and edited the manuscript. IY: conducted animal sampling and edited the manuscript. CS: conducted molecular analysis. EH: contributed with animal sampling and edited the manuscript. RT: conceptualised whole study and edited the manuscript. All approved manuscript.

## References

Ahmad, S. and Levy, L.S. (2010) ‘The Frequency of Occurrence and Nature of Recombinant Feline Leukemia Viruses in the Induction of Multicentric Lymphoma by Infection of the Domestic Cat with FeLV-945’. Virology 403 (2), 103–110

Biezus, G., Grima de Cristo, T., Bassi das Neves, G., da Silva Casa, M., Barros Brizola, P., Silvestre Sombrio, M., Miletti, L.C., and Assis Casagrande, R. (2023) ‘Phylogenetic Identification of Feline Leukemia Virus A and B in Cats with Progressive Infection Developing into Lymphoma and Leukemia’. Virus Research 329 (October 2022), 199093

Boomer, S., Eiden, M., Burns, C.C., and Overbaugh, J. (1997) ‘Three Distinct Envelope Domains, Variably Present in Subgroup B Feline Leukemia Virus Recombinants, Mediate Pit1 and Pit2 Receptor Recognition’. Journal of Virology 71 (11), 8116–8123

Brown, M.A., Cunningham, M.W., Roca, A.L., Troyer, J.L., Johnson, W.E., and Brien, S.J.O. (2008) ‘Characterization of FeLV from Florida Panthers’. Emerging Infectious Diseases [online] 14 (2). available from <https://www.cdc.gov/eid>

Burling, A.N., Levy, J.K., Scott, H.M., Crandall, M.M., Tucker, S.J., Wood, E.G., and Foster, J.D. (2017) ‘And Feline Immunodeficiency Virus Infection and Risk Factors for Seropositivity’. Small Animal 251 (2), 187–194

Cano-Ortiz, L., Tochetto, C., Roehe, P.M., Franco, A.C., and Junqueira, D.M. (2022) ‘Could Phylogenetic Analysis Be Used for Feline Leukemia Virus (FeLV) Classification?’ Viruses 14 (2)

Castillo-Aliaga, C., Blanchard, A.M., Castro-Seriche, S., Hidalgo-Hermoso, E., Jerez-Morales, A., Loose, M.W., and Tarlinton, R.E. (2023) ‘Comparison Between Sanger, Illumina and Nanopore Sequencing Evidencing Intra-Host Variation of Feline Leukemia Virus That Infects Domestic Cats’. BioRxiv [online] 2023.11.02.563952. available from <http://biorxiv.org/content/early/2023/11/02/2023.11.02.563952.abstract>

Castillo-Aliaga, C., Castro-Seriche, S., Jerez-Morales, A., and Tarlinton, R. (2024) ‘High Prevalence and Risk Factors of Feline Leukemia Virus Infection in Chilean Urban Cats (Felis Catus).’ Research in Veterinary Science [online] 180, 105403. available from <https://linkinghub.elsevier.com/retrieve/pii/S0034528824002704>

Chiu, E.S., Kraberger, S., Cunningham, M., Cusack, L., Roelke, M., and Vandewoude, S. (2019) ‘Multiple Introductions of Domestic Cat Feline Leukemia Virus in Endangered Florida Panthers’. Emerging Infectious Diseases 25 (1), 92–101

Choi, Y.R., Iturriaga, M.P., Nekouei, O., Tu, T., Van Brussel, K., Barrs, V.R., and Beatty, J.A. (2024) ‘Domestic Cat Hepadnavirus and Pathogenic Retroviruses; A Sero-Molecular Survey of Cats in Santiago, Chile’. Viruses 16 (1)

Coffin, J., Blomberg, J., Fan, H., Gifford, R., Hatziioannou, T., Lindemann, D., Mayer, J., Stoye, J., Tristem, M., and Johnson, W. (2021) ‘ICTV Virus Taxonomy Profile: Retroviridae 2021’. Journal of General Virology 102 (12), 1–2

Cunningham, M.W., Brown, M.A., Shindle, D.B., Terrell, S.P., Hayes, K.A., Ferree, B.C., McBride, R.T., Blankenship, E.L., Jansen, D., Citino, S.B., Roelke, M.E., Kiltie, R.A., Troyer, J.L., and O’Brien, S.J. (2008) ‘Epizootiology and Management of Feline Leukemia Virus in the Florida Puma’. Journal of Wildlife Diseases 44 (3), 537–552

Erbeck, K., Gagne, R.B., Kraberger, S., Chiu, E.S., Roelke-Parker, M., and VandeWoude, S. (2021) ‘Feline Leukemia Virus (FeLV) Endogenous and Exogenous Recombination Events Result in Multiple FeLV-B Subtypes during Natural Infection’. Journal of Virology 95 (18)

Faix, P.H., Feldman, S.A., Overbaugh, J., and Eiden, M. V. (2002) ‘Host Range and Receptor Binding Properties of Vectors Bearing Feline Leukemia Virus Subgroup B Envelopes Can Be Modulated by Envelope Sequences Outside of the Receptor Binding Domain’. Journal of Virology 76 (23), 12369–12375

Filoni, C., Helfer-Hungerbuehler, A.K., Catão-Dias, J.L., Marques, M.C., Torres, L.N., Reinacher, M., and Hofmann-Lehmann, R. (2017) ‘Putative Progressive and Abortive Feline Leukemia Virus Infection Outcomes in Captive Jaguarundis (Puma Yagouaroundi)’. Virology Journal 14 (1)

Gallina, L., Facile, V., Roda, N., Chiara, M., Alessia, S., Lorenza, T., Magliocca, M., Vasylyeva, K., Dondi, F., Balboni, A., and Battilani, M. (2024) ‘Molecular Investigation and Genetic Characterization of Feline Leukemia Virus (FeLV) in Cats Referred to a Veterinary Teaching Hospital in Northern Italy’. Veterinary Research Communications

Gregor, K.M., Mirolo, M., Brandes, F., Jesse, S.T., Kaiser, F., Verspohl, J., Wölfl, S., Osterhaus, A.D.M.E., Baumgärtner, W., Ludlow, M., and Beineke, A. (2024) ‘Fatal Feline Leukemia Virus-Associated Enteritis in a Wild Eurasian Lynx (Lynx Lynx) in Germany †’. Biology 13 (12)

Hardy, W.D., MacEwen, E.G., McClelland, A.J., Zuckerman, E.E., Myron, M., and Essex, E. (1976) ‘Biology of Feline Leukemia Virus in the Natural Environment’. Cancer Research 36 (February), 582–588

Hartmann, K. (2012) ‘Feline Leukemia Virus Infection’. in Infectious Diseases of the Dog and Cat. Fourth Edi. ed. by Greene, C.E. St Louis, Missouri: Elsevier, 108–136

Hartmann, K. and Hofmann-Lehmann, R. (2020) ‘What’s New in Feline Leukemia Virus Infection’. Veterinary Clinics of North America - Small Animal Practice 50 (5), 1013–1036

Helfer-Hungerbuehler, A.K., Widmer, S., Kessler, Y., Riond, B., Boretti, F.S., Grest, P., Lutz, H., and Hofmann-Lehmann, R. (2015) ‘Long-Term Follow up of Feline Leukemia Virus Infection and Characterization of Viral RNA Loads Using Molecular Methods in Tissues of Cats with Different Infection Outcomes’. Virus Research 197, 137–150

Lacerda, L.C., Silva, A.N., Freitas, J.S., Cruz, R.D.S., Said, R.A., and Munhoz, A.D. (2017) ‘Feline Immunodeficiency Virus and Feline Leukemia Virus: Frequency and Associated Factors in Cats in Northeastern Brazil’. Genetics and Molecular Research 16 (2)

Little, S., Levy, J., Hartmann, K., Hofmann-Lehmann, R., Hosie, M., Olah, G., and Denis, K.S. (2020) ‘2020 AAFP Feline Retrovirus Testing and Management Guidelines’. Journal of Feline Medicine and Surgery 22 (1), 5–30

Luaces, I., Doménech, A., García-Montijano, M., Collado, V.M., Sánchez, C., Tejerizo, J.G., Galka, M., Fernández, P., and Gómez-Lucía, E. (2008) ‘Detection of Feline Leukemia Virus in the Endangered Iberian Lynx (Lynx Pardinus)’. Journal of Veterinary Diagnostic Investigation 385 (2008), 381–385

Meli, M., Cattori, V., Martınez, F., López, G., Vargas, A., Palomares Francisco López-Bao, J. V., Hofmann-lehmann, R., and Lutz, H. (2010) Feline Leukemia Virus Infection□: A Threat for the Survival of the Critically Endangered Iberian Lynx (Lynx Pardinus). 134, 61–67

Miyake, A., Watanabe, S., Hiratsuka, T., Ito, J., Ngo, M.H., Makundi, I., Kawasaki, J., Endo, Y., Tsujimoto, H., and Nishigaki, K. (2016) ‘Novel Feline Leukemia Virus Interference Group Based on the Env Gene’. Journal of Virology 90 (9), 4832–4837

Mora, M., Napolitano, C., Ortega, R., Poulin, E., and Pizarro-Lucero, J. (2015) ‘Feline Immunodeficiency Virus and Feline Leukemia Virus Infection in Free-Ranging Guignas (Leopardus Guigna) and Sympatric Domestic Cats in Human Perturbed Landscapes on Chiloe Island, Chile’. Journal of Wildlife Diseases 51 (1), 199–208

Nájera, F., Grande-gómez, R., Peña, J., Vázquez, A., Palacios, M.J., Rueda, C., Corona-bravo, A.I., Zorrilla, I., Revuelta, L., Gil-molino, M., and Jiménez, J. (2021) ‘Disease Surveillance during the Reintroduction of the Iberian Lynx (Lynx Pardinus) in Southwestern Spain’. Animals 11 (2), 1–19

Nájera, F., López, G., del Rey-Wamba, T., Malik, R.A., Garrote, G., López-Parra, M., Fernández-Pena, L., García-Tardío, M., Arenas-Rojas, R., Simón, M.A., Zorrilla, I., Fernández, I., Alcaide, E.M., Ruiz, C., Revuelta, L., Salcedo, J., Hofmann-Lehmann, R., and Meli, M.L. (2024) ‘Long-Term Surveillance of the Feline Leukemia Virus in the Endangered Iberian Lynx (Lynx Pardinus) in Andalusia, Spain (2008–2021)’. Scientific Reports [online] 14 (1), 1–12. available from <10.1038/s41598-024-55847-3>

Ortega, C., Valencia, A.C., Duque-Valencia, J., and Ruiz-Saenz, J. (2020) ‘Prevalence and Genomic Diversity of Feline Leukemia Virus in Privately Owned and Shelter Cats in Aburrá Valley, Colombia’. Viruses 12 (4), 1–13

Ortega, R., Mena, J., Grecco, S., Pérez, R., Panzera, Y., Napolitano, C., Zegpi, N.A., Sandoval, A., Sandoval, D., González-Acuña, D., Cofré, S., Neira, V., and Castillo-Aliaga, C. (2020) ‘Domestic Dog Origin of Carnivore Protoparvovirus 1 Infection in a Rescued Free-Ranging Guiña (Leopardus Guigna) in Chile’. Transboundary and Emerging Diseases (August), 1–7

Palomares, F., Rodríguez, A., Revilla, E., López-Bao, J.V., and Calzada, J. (2010) ‘Assessment of the Conservation Efforts to Prevent Extinction of the Iberian Lynx’. Conservation Biology 25 (1), 4–8

Petch, R.J., Gagne, R.B., Chiu, E., Mankowski, C., Rudd, J., Roelke-Parker, M., Vickers, T.W., Logan, K.A., Alldredge, M., Clifford, D., Cunningham, M.W., Onorato, D., and VandeWoude, S. (2022) ‘Feline Leukemia Virus Frequently Spills Over from Domestic Cats to North American Pumas’. Journal of Virology

Polani, S., Roca, A.L., Rosensteel, B.B., Kolokotronis, S.O., and Bar-Gal, G.K. (2010) ‘Evolutionary Dynamics of Endogenous Feline Leukemia Virus Proliferation among Species of the Domestic Cat Lineage’. Virology [online] 405 (2), 397–407. available from <10.1016/j.virol.2010.06.010>

Powers, J.A., Chiu, E.S., Kraberger, S.J., Roelke-parker, M., Lowery, I., Erbeck, K., Troyer, R., Carver, S., and VandeWoude, S. (2018) ‘Crossm Feline Leukemia Virus (FeLV) Disease Outcomes in a Domestic Cat Breeding Colony□: Relationship to Endogenous FeLV and Other Chronic Viral Infections’. Journal of Virology 92 (18), 1–16

Rather, M.A., Hassan, A., Aman, M., Gul, I., Mir, A.H., Potdar, V., Koul, P.A., Ahmad, S.M., Ganai, N.A., Shah, R.A., Chikan, N.A., Abdul-Careem, M.F., and Shabir, N. (2025) ‘Molecular and Ecological Determinants of Mammalian Adaptability in Avian Influenza Virus’. in Infection. Springer Science and Business Media Deutschland GmbH

Rohn, J.L., Linenberger, M.L., Hoover, E.A., and Overbaugh, J. (1994) ‘Evolution of Feline Leukemia Virus Variant Genomes with Insertions, Deletions, and Defective Envelope Genes in Infected Cats with Tumors’. Journal of Virology 68 (4), 2458–2467

Sacristán, I., Acuña, F., Aguilar, E., García, S., José López, M., Cabello, J., Hidalgo-Hermoso, E., Sanderson, J., Terio, K.A., Barrs, V., Beatty, J., Johnson, W.E., Millán, J., Poulin, E., and Napolitano, C. (2021a) ‘Cross-Species Transmission of Retroviruses among Domestic and Wild Felids in Human-Occupied Landscapes in Chile Asdasdasd’. Evolutionary Applications 14 (4), 1070–1082

Sacristán, I., Esperón, F., Pérez, R., Acuña, F., Aguilar, E., García, S., López, M.J., Neves, E., Cabello, J., Hidalgo-Hermoso, E., Terio, K.A., Millán, J., Poulin, E., and Napolitano, C. (2021b) ‘Epidemiology and Molecular Characterization of Carnivore Protoparvovirus-1 Infection in the Wild Felid Leopardus Guigna in Chile’. Transboundary and Emerging Diseases 68 (6), 3335–3348

Stewart, H., Jarrett, O., Hosie, M.J., and Willett, B.J. (2013) ‘Complete Genome Sequences of Two Feline Leukemia Virus Subgroup B Isolates with Novel Recombination Sites’. Genome Announcements 1 (1), 1–2

Studer, N., Lutz, H., Saegerman, C., Gönczi, E., Meli, M.L., Boo, G., Hartmann, K., Hosie, M.J., Moestl, K., Tasker, S., Belák, S., Lloret, A., Boucraut-Baralon, C., Egberink, H.F., Pennisi, M.-G., Truyen, U., Frymus, T., Thiry, E., Marsilio, F., Addie, D., Hochleithner, M., Tkalec, F., Vizi, Z., Brunetti, A., Georgiev, B., Ludwig-Begall, L.F., Tschuor, F., Mooney, C.T., Eliasson, C., Orro, J., Johansen, H., Juuti, K., Krampl, I., Kovalenko, K., Šengaut, J., Sobral, C., Borska, P., Kovaříková, S., and Hofmann-Lehmann, R. (2019) ‘Pan-European Study on the Prevalence of the Feline Leukaemia Virus Infection-Reported by the European Advisory Board on Cat Diseases (ABCD Europe)’. Viruses 11, 993

Torres, A.N., Mathiason, C.K., and Hoover, E.A. (2005) ‘Re-Examination of Feline Leukemia Virus: Host Relationships Using Real-Time PCR’. Virology 332 (1), 272–283

Watanabe, S., Kawamura, M., Odahara, Y., Anai, Y., Ochi, H., Nakagawa, S., Endo, Y., Tsujimoto, H., and Nishigaki, K. (2013) ‘Phylogenetic and Structural Diversity in the Feline’. PLoS ONE 8 (4), 1–16

Willett, B.J. and Hosie, M.J. (2013) ‘Feline Leukaemia Virus: Half a Century since Its Discovery’. Veterinary Journal 195 (1), 16–23

